# Abundance-activity decoupling in sulfur-cycling bacteria reflects viral infection types in meromictic lakes

**DOI:** 10.64898/2026.03.13.711613

**Authors:** Jordan R. Walker, Natascha S. Varona, Bailey A. Wallace, Abelardo Aguilar, Molly D. O’Beirne, Josef P. Werne, Antoni Luque, William P. Gilhooly, Alice Bosco-Santos, Cynthia B. Silveira

## Abstract

Meromictic lakes serve as analogs for redox-stratified ancient oceans with well-mixed surface waters and anoxic bottoms. Purple and green sulfur bacteria (PSB, GSB) dominate the anoxic zones where light penetrates, and their biosignatures can be used to guide interpretations of geologic records. However, PSB and GSB biosignatures do not directly reflect the microbial community composition of modern lakes, posing a challenge for their interpretation. Here, we investigate this decoupling by integrating metagenomics, metatranscriptomics, and metaHi-C virus–host linkages with the geochemical profiles of three meromictic lakes. In the phototrophic microbial plates, PSBs’ transcriptional activity far exceeded their abundance (73% of the total microbial community activity versus 30% of the abundance), while GSBs displayed the opposite pattern. Concurrently, PSBs were exclusively associated with temperate viruses, while GSBs were targeted by lytic infections. Sulfate-reducing bacteria (SRB) and viruses encoding genes for sulfate reduction were most active where sulfide concentration was lowest. These results reveal that viral replication strategies are associated with the decoupling between abundance and activity in anoxygenic phototrophs and sulfate reducers. These relationships could accelerate sulfur regeneration, contribute to sustaining phototrophy, and ultimately reflect in the lake’s bulk biosignatures.

## Introduction

In the Precambrian, prior to the oxygenation of Earth’s atmosphere (> 2.4 Ga), anoxygenic photosynthetic bacteria were major contributors to marine primary production [1,2]. During the Paleozoic, their contribution became context-dependent, expanding during transient episodes of oceanic oxygen limitation, particularly in the Early Paleozoic (540-470 Ma) and Devonian (420-360 Ma) oceans [3–7]. Today, anaerobic photoautotrophs persist as purple and green sulfur bacteria (PSB and GSB), which can dominate illuminated anoxic environments such as meromictic lakes, euxinic and ferruginous basins, and permanently stratified coastal waters [8–12].

Biosignatures (i.e., metabolic outputs) produced by PSB and GSB in modern redox-stratified systems provide a powerful window into anoxygenic primary production [13–15]. In this context, meromictic lakes have been extensively used to investigate past anoxic environments, to inform interpretations of anoxic settings beyond Earth, and to anticipate future ocean deoxygenation [5,9,16–19]. These lacustrine systems are stratified into three distinct layers: an oxygenated seasonally mixed surface layer (mixolimnion), an intermediate chemocline characterized by steep redox and chemical gradients, and a stable anoxic bottom layer (monimolimnion). The chemocline, where light penetrates sulfide-rich waters, provides a unique niche for anoxygenic photoautotrophs to thrive. As such, meromictic lakes can be used to investigate the processes that control biosignature production by PSB and GSB, and their preservation in the geological record.

In illuminated anoxic environments, PSB and GSB typically oxidize reduced sulfur compounds, most commonly sulfide, to fuel photosynthesis [11,20]. In these settings, they are metabolically coupled to sulfate-reducing bacteria (SRB) through microbial sulfur cycling. Sulfide produced by SRB helps to fuel anoxygenic photosynthesis by PSB and GSB, while organic carbon and intermediate sulfur compounds produced by phototrophic sulfur bacteria support heterotrophic metabolisms, including sulfate reduction [21,22]. This tight coupling establishes steep redox gradients, linking sulfur oxidation and reduction within chemoclines [23,24]. As sulfur cycling bacteria, SRB also leave diagnostic biosignatures in the geological record, most notably distinct sulfur isotopic fractionations that extend back more than 3 billion years [25–27].

Despite the extensive use of PSB and GSB biosignatures to reconstruct anoxic environments, a growing body of evidence indicates that biomarker abundance and carbon and sulfur isotopic ratios are often decoupled from the abundance and composition of sulfur bacterial communities [5,28,29]. Bottom-up controls, such as light and sulfide availability, have been insufficient to explain this decoupling [17,23,30–33]. One potential, yet underexplored, explanation for this decoupling is viral modulation of host metabolism. Viruses of bacteria can establish lytic (virulent) infections that kill the host cell, or different types of long-term interactions, such as chronic and lysogenic infections, where the virus replicates without killing the host or integrates into the bacterial genome, respectively. In all cases, viruses often encode metabolic genes that can be expressed during infections, thereby altering bacterial cell physiology [34–37]. Viral infection can also expand host metabolic capabilities via lateral gene transfers [38,39] and changes in overall gene expression profiles [40–42]. It is, therefore, possible that viral infections represent a mechanism causing a decoupling between abundance and metabolism, and consequently, biosignatures of anoxygenic phototrophs in anoxic and sulfidic lakes. Recent observations supporting this possibility include the identification of viruses encoding metabolic genes capable of influencing carbon fixation, sulfur cycling, and pigment production in meromictic lake waters and sediments, and the transitions in viral replication strategy indicated by viral abundance, virus-to-microbe ratios, and viral transcriptional activity in these lakes [18,19,43].

While precedent exists supporting the role of viruses in altering microbial community structure and metabolic function in meromictic systems, viral interactions with sulfur-metabolizing bacteria remain poorly constrained. In particular, the relative importance of lytic versus lysogenic infection strategies, the specificity of virus-host associations, and the extent to which viral-encoded genes influence sulfur cycling and carbon fixation are not well understood. To address these gaps, here we integrate metatranscriptomic, metagenomic, and proximity-ligation (metaHi-C) approaches to study the relationship between the activity of sulfur-metabolizing bacteria and the viral replication and gene expression dynamics in three meromictic lakes spanning from trace (0.006 mM) to high (25.389 mM) sulfide concentration systems.

## Materials & Methods

### Sampling and geochemical data

The three study lakes, Mahoney Lake (British Columbia, Canada), Poison Lake (Washington, USA), and Lime Blue Lake (Washington, USA), were sampled between July 19^th^ and July 25^th^, 2023. Briefly, water column temperature, dissolved oxygen (DO), specific conductivity, and pH were measured using a YSI ProDSS multiparameter probe (YSI Incorporated, Yellow Springs, OH, USA) and photosynthetically active radiation (PAR) was measured using a spherical underwater quantum sensor coupled to a data logger (LI-193 and LI-1500; LI-COR Environmental, Lincoln, NE, USA) (Supplemental Table S1). Water samples for geochemical and microbiological analyses were collected from three discrete depths in five replicates (surface, microbial plate, and bottom) using a peristaltic pump (Electra; Proactive Environmental Products, Bradenton, FL, USA) coupled to lateral intake, minimizing vertical mixing (Supplemental Table S2). Sampling depths were: 3.0 m, 8.78 m, and 13.0 m for Mahoney Lake (MAH), 3.0 m, 6.7 m, and 7.5 m for Poison Lake (POI), and 3.0 m, 12.0 m, and 13.7 m for Lime Blue Lake (LB). Dissolved sulfide concentrations were measured in 5 mL samples obtained from the chemoclines of the three lakes, fixed with 0.05 M Zn-acetate (Zn(CH3COO)2•2H2O) solution at a 1:2 ratio immediately after sampling, and quantified photometrically using the methylene blue method [44].

Samples for total nucleic acid extractions were collected in replicate using 0.22 μm PES Sterivex filters (Millipore-Sigma, Burlington, MA). Five replicate samples were collected per lake and depth for DNA extraction and five for RNA extraction, for a total of 90 Sterivex filters. Filtered volumes varied with microbial density at each lake depth due to clogging; volumes are reported in Supplemental Table S2. Immediately after collection, 1 mL of RNAlater (Invitrogen, Carlsbad, CA) was added to Sterivex filters for metatranscriptomic analyses, flash-frozen in liquid nitrogen and stored in dry shippers until their return to the University of Miami laboratory, where they were stored at -80°C. An additional 3 mL of microbial plate water from each lake was filtered through 0.2 μm, 25 mm cellulose nitrate filters (cat #10400106, Whatman, Milwaukee, USA) for metaHi-C cross-linking and library preparation. These filters were also flash-frozen in liquid nitrogen immediately after collection and stored at -80°C until processing.

### Library preparation and sequencing

Total DNA was extracted from the 45 metagenomic samples using a modified Nucleospin Tissue Kit (Macherey-Nagel, Duren Germany). The protocol was modified by adding 360 μL of T1 buffer and 50 μL of Proteinase K directly to the Sterivex filters, followed by overnight incubation at 55°C. The following day, 400 μL of B3 buffer was added to each Sterivex, and samples were incubated at 70°C for 30 minutes. The lysate was recovered from the Sterivex filters and mixed with 420 μL of 100% ethanol. The resulting mixture was loaded onto silica spin columns in 600 μL increments and centrifuged at 11,000 x g for 1 minute. This step was repeated until the entire sample volume had passed through the column. Washing steps were performed according to the manufacturer’s instructions. DNA was eluted in 50-100μL of molecular-grade water. DNA concentrations were quantified using a Qubit 2.0 Fluorometer (Life Technologies, Carlsbad, CA, USA). For samples with low DNA yield or large elution volumes, DNA was concentrated by ethanol precipitation through the addition of 0.1× the elution volume of 3 M sodium acetate, 2.5× the elution volume of ice cold 100% ethanol, and 1 μL of glycogen, followed by incubation at -20°C for 3 hours. Samples were centrifuged at 4°C for 30 min at 21,130 x g, after which the supernatant was removed. The DNA pellet was washed with 500 μL of 75% ethanol and centrifuged at 4°C for 10 min at 21,130 x g. This wash step was repeated, after which the pellet was briefly spin-dried (10 seconds) and resuspended in 50 μL of molecular-grade water. DNA extracts were stored at -80°C and shipped to the DOE Joint Genome Institute (JGI) for sequencing on an Illumina NovaseqX, yielding a total of 5,668,261,550 paired-end 2×151bp reads (Supplemental Table S3).

Metatranscriptomes were sequenced and published in [19], and were re-analyzed here in combination with newly generated metagenomes and metaHi-C data. Briefly, metatranscriptomic samples were processed using a modified DNeasy PowerWater Sterivex kit (Qiagen, Germantown, MD, USA) following Krausfeldt, et al. [45]. Fourteen samples failed RNA extraction; for two of these, RNA was extracted from unfiltered flash-frozen water samples using a chloroform/TRIzol reagent (Invitrogen, Waltham, MA, USA). Following treatment with TURBO DNase (Thermo Fisher Scientific, Waltham, MA, USA) and rRNA depletion using QIAseq® FastSelect™ − rRNA 5S/16S/23S kit (Qiagen, Germantown, MD, USA), cDNA libraries were constructed with NEBNext Ultra II RNA Library Preparation Kit for Illumina and sequenced using Illumina NovaSeq platform, generating a total of 766,883,916 paired-end 2 x 150bp reads (Supplemental Table S4).

Hi-C libraries were prepared using the ProxiMeta™ Hi-C v4.0 Kit (Phase Genomics, Seattle, USA). Filters were cross-linked with formaldehyde and digested with the restriction enzymes Sau3AI and MlucI. DNA fragments were proximity ligated with biotinylated nucleotides and purified with streptavidin beads prior to library preparation using an Illumina-compatible kit. Hi-C libraries were sequenced on an Illumina Novaseq, producing ∼160 M paired-end 2 x 150 bp reads. Hi-C library preparation failed for Lime Blue Lake microbial plate samples; therefore, only Poison Lake and Mahoney Lake chemocline samples yielded Hi-C data.

### Genome binning

Metagenomic reads were processed according to JGI’s standard metagenome workflow [46]. This pipeline includes quality control filtering (bbdusk.sh parameters maq=3.0,0 trimq=0.0 qtrim=r maxns=3 minlen=51 minlenfraction=0.33 k=31 hdist=1), adapter and artifact trimming (bbduk.sh parameters ktrim=r minlen=51 minlenfraction=0.33 mink=11 tbo tpe rcomp=f k=23 hdist=1 hdist2=1 ftm=5 pratio=G,C plen=20), masking of common laboratory and microbial contaminants (bbmap.sh parameters deterministic quickmatch k=13 idtag=t qtrim=rl trimq=10 untrim minid=.95 idfilter=.95 maxindel=3 minhits=2 bw=12 bwr=0.16 maxsites2=10 tipsearch=0, and read error correction to attempt to correct low coverage sequencing errors (bbcms.sh parameters mincount=2 and highcountfraction=0.6) using the BBTools Software package version 39.03 [47] (Supplemental Table S3). Assembly of contigs and scaffolds was performed with metaSPAdes (version 3.15.2; parameters -m 2000 --only-assembler -k 33,55,77,99,127 --meta) (Supplemental Table S3), reads were mapped using BBMap (parameters ambiguous=random), and scaffolds were binned into metagenome assembled genomes (MAGs) with MetaBat2 (version 2.15) using standard parameters [47–49]. MAG completeness and contamination was estimated using CheckM 2 (version 2.15) [50]. MAGs with more than 50% completion and containing less than 10% contamination were retained for further analysis (N = 1,430). Metatranscriptomic reads were quality-controlled and assembled according to [19] using BBDuk (version 38) and metaSPAdes (version 3.15.5), respectively. Hi-C reads were preprocessed to cut reads at restriction enzyme sites using hicstuff’s (version 3.2.4) digest function [51]. Hi-C reads were combined with their corresponding scaffolds from the metagenomes described above and binned using Metator version 1.3.10 [52]. This resulted in 5 separate Hi-C binning sets for both Mahoney and Poison microbial plates, one for each replicate metagenome. Viral sequences (identified in the following section) were removed to yield the final set of prokaryotic MAGs for downstream analyses. CRISPR spacers were predicted from these MAGs using MinCED version 0.4.2 [53]. Metagenome and Hi-C derived MAGs were dereplicated using dRep version 3.5.0 with standard setting [54]. Open reading frames were predicted with Prodigal (version 2.6.3), taxonomic assignment with GTDB-tk (version 2.4.1), anti-phage systems with defensefinder (version 2.0.1) and protein annotation of translated amino acid sequences with MetaCerberus (version 1.2.1) [55–58] (Supplemental Table S3).

### Identification of viral genomes

Viral sequences were identified among the metagenomic scaffolds, metatranscriptomic contigs, and quality-filtered metagenomic and Hi-C MAGs using geNomad (version 1.8.0) and VIBRANT (version 1.2.1) [59,60]. Predicted viral sequences longer than 2.5 Kbp were clustered into viral Operational Taxonomic Units (vOTUs) using the anicalc and aniclust scripts in CheckV (version 1.0.3) with an average nucleotide identity threshold of 95% and a minimum coverage of 85%, following MIUVIG standards [61,62]. CheckV was used to exclude representative vOTU sequences (hereafter simply referred to as vOTUs) with no viral hallmark genes and remove host regions flanking vOTUs identified as prophages. MetaCerberus with prodigal-gv was used for vOTU gene annotation [56,57,59]. Putative temperate viral genomes were identified among vOTUs using VIBRANT parameters for prophage identification and keywords in the MetaCerberus annotations, including integrase, recombinase, transposase, excisionase, Cro/C1, phage (anti)repressor, PHROG or VOG-derived ParAB, and phage regulatory protein CII. Integrated prophages were identified using a combination of 3 different methods: 1) vOTUs that contained member sequences identified as prophages with host flanking regions during viral identification by geNomad or VIBRANT, 2) identification of prophages using CheckV, and 3) BLASTn (version 2.16.0) alignment of pre-dereplicated MAGs to vOTUs with at least 95% identity, 100% coverage of the vOTU, and 500 bp flanking regions on each end of each viral sequence [63]. Putative viral hosts were identified in four ways: 1) alignment of CRISPR spacers from dereplicated MAGs to vOTUs with 100% coverage, less than two mismatches, and a minimum length of 20 bp, 2) prophages integrated into MAGs, 3) host prediction using iPHoP with a minimum confidence score of 0.95 and 4) sequences identified as viral using geNomad or VIBRANT within Hi-C MAGs created using Metator [63,64]. All unique virus-host interactions observed were compiled into a bipartite matrix and visualized in Cytoscape, where host were classified at the order level and viruses were classified according to lifestyle [65]. We employed the use of Virsosrter2 to further inspect PSB MAGs for potential prophages that may have been missed in the pipeline utilized for metagenomes, metatranscriptomes, and HiC MAGs [66].

### Calculation of relative abundance and activity

Metagenomic and metatranscriptomic reads were mapped to dereplicated MAGs and vOTUs using BBMap with standard settings. Resulting SAM files were converted to BAM format, sorted, and indexed using samtools (version 1.21) [67]. The number of reads or transcripts mapping to each contig was quantified using CoverM (version 0.7.0) [68]. Fractional abundances and transcription of MAGs and vOTUs were calculated using the equation:

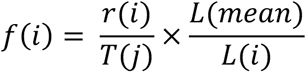

where r(i) is the number of reads/transcripts mapped to a MAG or vOTU, T(j) is the total number of reads/transcripts in a metagenome/metatranscriptome, L(mean) is the mean contig or vOTU length, and L(i) is the length of a contig or vOTU. The relative abundances and transcription were calculated as f(i) divided by the sum of all f(i) within a metagenome/metatranscriptome.

Statistical analysis and visualization were done in RStudio (version 2025.05.0+496) and R (version 4.4.2) [69,70]. Statistical tests were performed using the package rstatix (version 0.7.2) [71]. Data visualization was performed using ggplot2 (version 3.5.2), ggpubr (version 0.6.1), and ggsci (version 3.2.0) [72–74].

## Results

### Environmental gradients and microbial plate characterization

Dissolved oxygen concentrations peaked at ∼5 meters depth in all three lakes, followed by a sharp transition to anoxia at ∼7-8 meters at Mahoney and Poison Lakes and a more gradual decline to anoxic conditions at ∼10 meters at Lime Blue Lake (Figure 1). Photosynthetically active radiation (PAR) decreased steadily with depth and approached zero at the microbial plate sampling depths in all lakes (Figure 1A). Depleted dissolved oxygen concentrations, lower PAR values, and visible color changes observed on 0.45 µm filters were used to define the sampling depth of the microbial plate. Based on these criteria the microbial plates were defined at 8.78 m in Mahoney Lake, 6.7 m in Poison Lake, and 12 m in Lime Blue Lake. Sulfide concentration varied by several orders of magnitude across the sampled microbial plates, ranging from 0.006 mM in Lime Blue Lake to 0.831 mM in Poison Lake, and reaching a maximum of 25.389 mM in Mahoney Lake.

**Figure 1.**
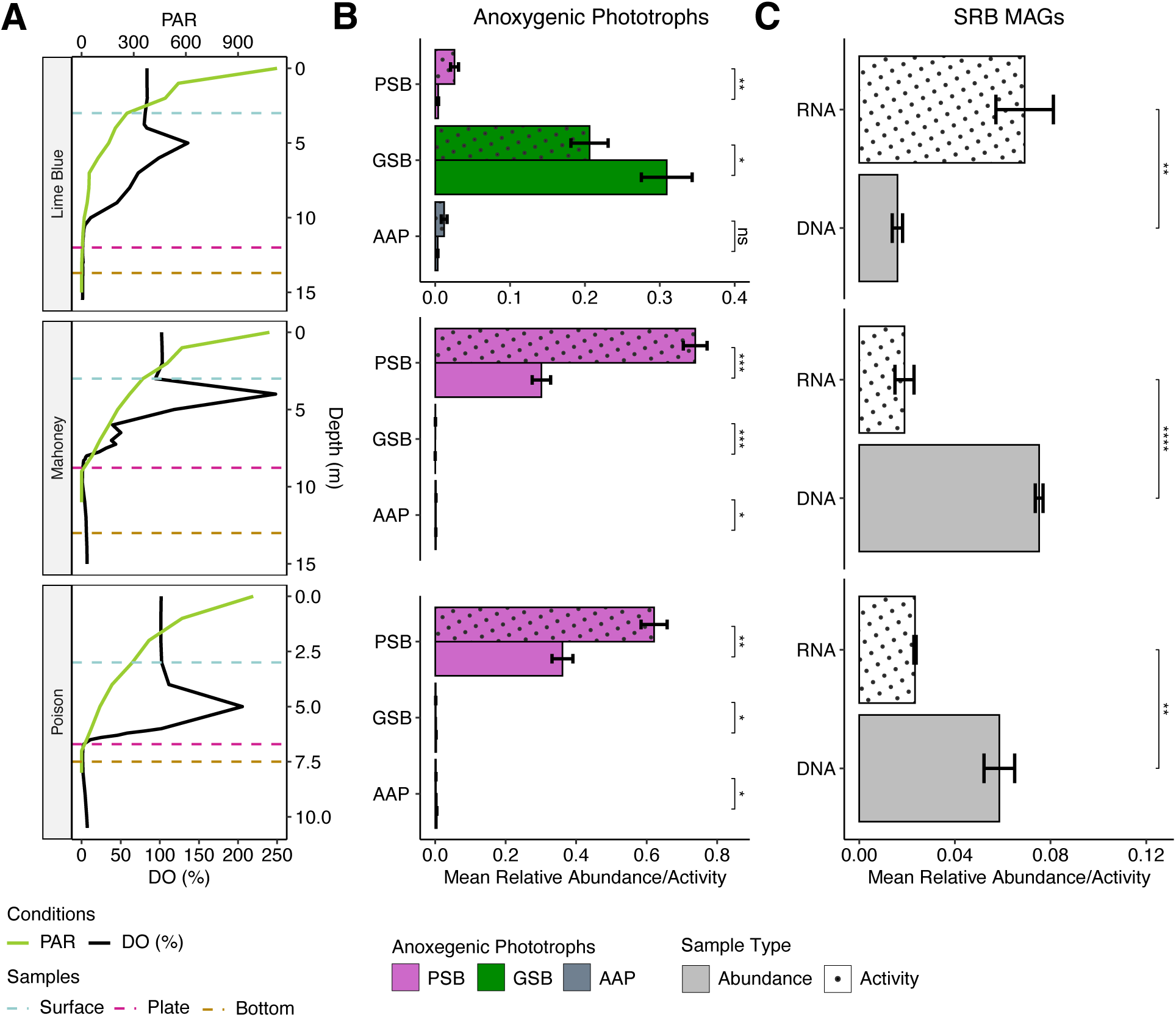
Depth profiles and distribution of anoxygenic phototrophs and sulfate reducers across study lakes. A) Vertical profiles of dissolved oxygen (%; black lines) and photosynthetically active radiation (PAR, green lines) for each lake (lake names shown at left). Dotted horizontal lines indicate the depths at which microbial communities were sampled, including the surface (blue), the microbial plate (purple), and the bottom (gold) layers. B-C). Abundance (solid) and activity (dotted) of anoxygenic phototrophs (B) and sulfate-reducing bacteria (C) in microbial plate samples. Note the different y-axis scales between panels B and C, where panel B’s scale is ∼7× larger than that of panel C. Anoxygenic phototrophs are grouped as PSBs (purple), GSBs (green), or aerobic anoxygenic phototrophs (gray). Results of Tukey’s tests comparing abundance to activity are denoted by ns = not significant, * = 0.05, ** = 0.01, *** = 0.001, **** = 0.0001.

### Abundance and transcriptional activity of anoxygenic phototrophs

Assembly and binning of metagenomic and Hi-C reads resulted in 580 high or medium quality, unique MAGs. Of these, 513 were classified as Bacteria and 67 as Archaea. Thirteen MAGs were classified as potential anoxygenic phototrophs based on the presence of genes encoding anoxygenic photosynthetic reaction centers, including *pufLM* and photosystem P840. Two anoxygenic phototroph MAGs were classified as PSBs within the order *Chromatiales*, genera *Chromatium* and *Thiohalocapsa*. The *Thiohalocapsa* MAG was the most abundant MAG in the microbial plates of Mahoney and Poison Lakes, accounting for 30% and 36% of the microbial community, respectively, and 73% and 62% of the community transcriptional activity (herein after referred to as activity) (Figure 1B). In contrast, the *Chromatium* MAG accounted for less than 1% of the community and activity in all samples, except in Lime Blue’s microbial plate, where it accounted for 0.03% of the abundance but its activity reached 2%. Across all three microbial plates, PSB exhibited significantly higher activity than what would be predicted from a direct comparison with their abundance (Tukey’s Test, α < 0.05).

Four anoxygenic phototrophic MAGs were classified as GSBs, including three belonging to the genus *Chlorobium* and one *Prosthecochloris*. One *Chlorobium* MAG dominated the Lime Blue bacterial plate, accounting for 29% of the community and 20% of the activity (Figure 1B). The remaining GSB MAGs each contributed < 2% of the community and less than 1% of the activity on the Lime Blue microbial plate. The GSBs’ activity was significantly lower than what would be predicted by their abundance (Tukey’s Test, α < 0.05) in all the microbial plates (Figure 1B).

The remaining seven anoxygenic phototrophic MAGs belonged to the orders *Burkholderiales*, *Caulobacterales*, *Rhodobacterales*, *Sphingomonadales*, and *Chthoniobacterales*. Except for *Chthoniobacterales*, all these orders contain known aerobic anoxygenic phototrophs. Consistent with their aerobic nature, aerobic anoxygenic phototrophs had the highest abundances and activities in Mahoney and Poison surface waters (Supplementary Figure 1A). Together, aerobic anoxygenic phototrophs accounted for less than 1% of community composition and activity in all bacterial plates except for Lime Blue, where they accounted for 1% of the activity (Figure 1B).

MAGs classified as sulfate-reducing bacteria (SRB) (orders Desulfobacterales (n=11), Desulfobulbales (n=7), and Desulfatiglandales (n=10) made up a much smaller fraction of the microbial community abundance and activity compared to the phototrophs (Figure 1B-C, note differences in scale). SRBs were significantly more active than their abundance would predict in the Lime Blue bacterial plate (7% and 1.6%, respectively), while the opposite was observed in Mahoney and Poison Lakes (Tukey’s Test, α < 0.05 for all comparisons, Figure 1). SRB were most abundant in bottom water samples, where abundance and activity were often coupled, except for Lime Blue, where SRBs showed higher activity than abundance (Supplementary Figure 1B).

### Viral community abundance and transcriptional activity

A total of 203,946 unique sequences were classified as putative viral genomes or genome fragments greater than 2.5 kbp (189,27 by geNomad and 98,544 by VIBRANT, with 83,882 identified by both viral prediction tools). Clustering and quality control of the viral contigs resulted in 44,392 viral operational taxonomic units (vOTUs; hereafter referred to as viruses), 4,269 of which were classified as temperate, and of those, 1,895 were identified as integrated prophages. Lytic virus activity was significantly higher than their abundance in all samples except for the microbial plates of Mahoney and Poison (Tukey’s Test, α < 0.05, Figure 2A, Supplementary Figure 2A). Temperate viral activity was highest at these two microbial plates (35% and 15% of the viral activity, respectively, Figure 2B, Supplementary Figure 2B). Temperate virus activity was only higher than abundance in the plates of Poison and Mahoney Lakes (Tukey’s test, α < 0.05 except for Mahoney, where the test was not significant).

**Figure 2.**
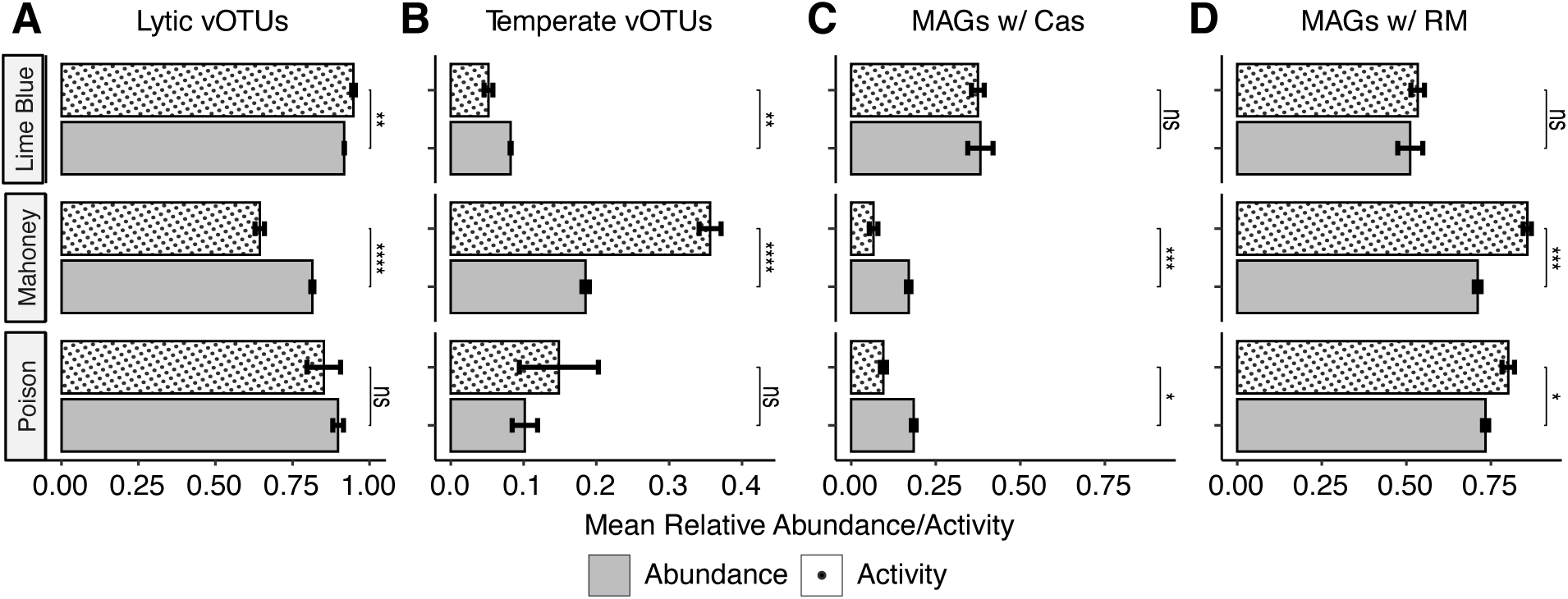
Abundance and activity of lytic and temperate viruses and hosts with defense systems in the microbial plates. Abundance is indicated by solid gray bars and activity by dotted, white bars. A) Viruses classified as lytic, (B) viruses classified as temperate, including prophages, (C) MAGs with genes involved in Cas defense systems, and (D) MAGs with genes annotated as RM systems. Results of Tukey’s tests comparing abundance to activity are denoted by ns = not significant, * = 0.05, ** = 0.01, *** = 0.001, **** = 0.0001.

Seventy-four percent of prokaryotic MAGs in the dataset carried phage defense systems, including all the PSB and GSB MAGs. MAGs encoding Cas systems were more abundant in the Lime Blue bacterial plate, accounting for 38 and 37% of the community abundance and activity, respectively (Figure 2C). In Mahoney and Poison Lakes, MAGs with Cas systems were less abundant (18 and 17%, respectively) and displayed lower activity relative to their abundances (Figure 2C). MAGs containing restriction modification (RM) systems showed the opposite trend, with a higher overall abundance and activity in Poison and Mahoney relative to Lime Blue, and higher activity relative to abundances in Mahoney and Poison Lakes (Tukey’s Test, α < 0.05, Figure 2D). The only other sample in which MAGs with Cas and RM systems displayed high activity relative to abundance was the bottom layer of Poison Lake (Supplementary Figure 2D).

### Virus-host infection network

Four virus-host linkage methods (MetaHi-C, CRISPR-spacer matches, prophage-matches, and iPHoP) identified 1,013 unique associations between viruses and MAGs (Supplementary Figure 3A). This network consisted of 970 viruses and 247 MAGs, where 43 viruses interacted with more than one MAG (Supplementary Figure 3). Hi-C links made up 63.6% of the total interactions, even though Hi-C data came from only two samples, Mahoney and Poison microbial plates (Supplementary Figure 3B-C). To investigate virus-host specificity we analyzed the structure of the network and found it to be highly fragmented, displaying 221 connecting groups with an average of 1.7 neighbors (Supplementary Figure 3). This fragmentation likely reflects the network’s representation of all virus-host interactions across different sample types (surface, microbial plate, and bottom waters), as well as false-negative linkages.

In the bacterial plates of Mahoney and Poison lakes, we detect that only temperate viruses interacted with the most active and abundant PSB genus, *Thiohalocapsa* (Figure 3A, Supplementary Figure 3). By contrast, all GSBs, including the most dominant in Lime Blue, *Chlorobium*, were associated exclusively with putatively lytic viruses (Figure 3A, Supplementary Figure 3). Virus-to-host ratios (VHR) were calculated from metagenomes and transcriptomes to indicate virus replication and activity relative to their hosts (Figure 3). Most of the virus–host pairs outside of the interquartile ranges (dotted lines, n=797, Figure 3B) fell into the upper right quadrant (high abundance and activity relative to host, n=351, 44%) or the lower left quadrant (low abundance and activity relative to host, n=341, 43%). Temperate viruses were enriched in the lower left quadrant (37%) compared to the upper right quadrant (18%). PSB-virus pairs fell exclusively within the interquartile range or in the lower-left quadrant, consistent with temperate viral infections characterized by relatively low lytic viral activity. On the other hand, GSB-virus pairs showed distinct patterns: the most dominant GSB in the chemocline of Lime Blue exhibited the lowest VHR abundance and activity levels in the dataset (Figure 3). The low activity and abundance (<1×10^-5^) of these GSB viruses, coupled with high host abundance and activity, resulted in the lowest VHRs in the dataset (Figure 3A). Another less common GSB, 1.5% abundance in the bottom and 0.3% in the chemocline, was associated with one of the most abundant and active viruses, leading to its inclusion in the upper right quadrant (Figure 3A). SRBs were associated with a diversity of lytic and temperate viruses, resulting in a broad range of VHRs, indicating high variability in viral activity and replication.

**Figure 3.**
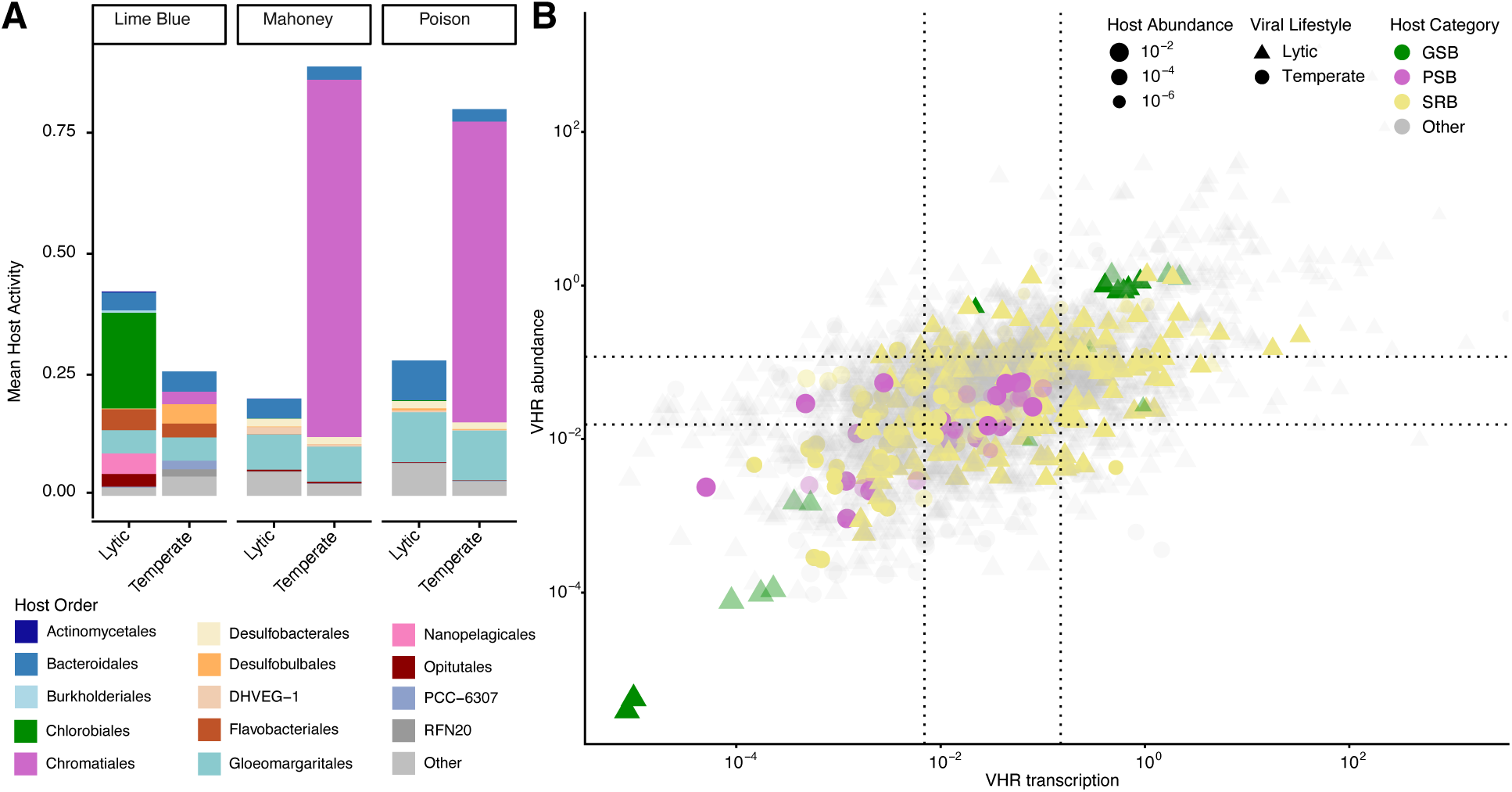
Virus-host dynamics of meromictic lakes. (A) The mean activity of hosts associated with lytic and temperate viruses. (B) Relationship between the virus-to-host ratio (VHR) calculated from abundances (Y-axis) and activity (X-axis) across all samples. Transparent points include all other bacteria and PSB, GSB, or SRB virus-host associations on the surface or bottom of the lakes. Opaque points represent viruses with connections to PSB (purple), GSB (green), or SRB (yellow) in the microbial plates. Triangles represent putatively lytic vOTUs and circles indicate temperate vOTUs. Point sizes are scaled to the log-transformed host abundance. Colors of bars represent the taxonomic classification of the host at the order level.

### Metabolic genes encoded by viruses

We searched for KEGG metabolic pathways most likely to have a direct effect on biosignature and biomarker production, including carbon fixation, glycolysis and gluconeogenesis, photosynthesis, antenna proteins, porphyrin metabolism, sulfur metabolism, and sulfur relay system. We found that there were more active viruses and viruses containing metabolic genes of interest in the Lime Blue bacterial plate (n=12,704; viruses with metabolic genes n=247) compared to Mahoney (n=3,319; viruses with metabolic genes n=84) or Poison (n=4,066; viruses with metabolic genes n=65, Figure 4A). In Mahoney and Poison lakes, approximately 50% and 51% (n=42, n=33), respectively, of the viruses carrying these metabolic genes were temperate, whereas in Lime Blue, only 31% (n=76) were found to be temperate.

**Figure 4.**
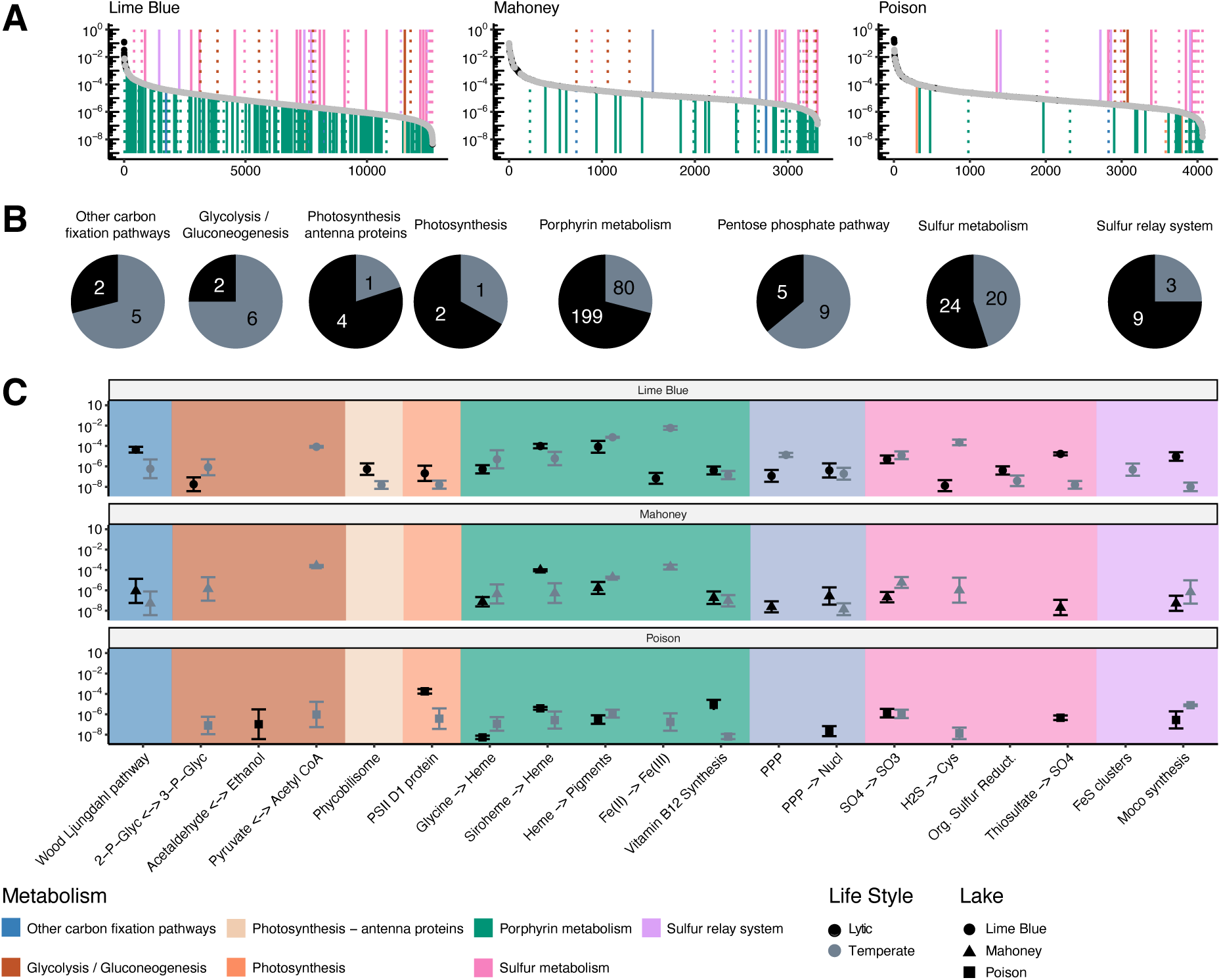
Distribution of viruses carrying metabolic genes related to biosignatures. A) Rank activity of viruses, where temperate viruses are denoted with gray dots. Vertical lines identify viruses carrying metabolic genes, colored according to their KEGG metabolic pathways, and line type denoting lifestyle: lytic (solid) or temperate (dotted). B) The counts of putatively lytic (black) and temperate (gray) viruses carrying metabolic genes within the pathways related to biosignatures across all lakes. C) Total activity of all viruses carrying genes involved in chemical reactions or pathways that could influence biosignatures. Prior to summation of genes within a pathway, activity was log(a + x) transformed, where a is the lowest, non-zero activity level for any one gene across all lakes divided by 10 (a = 3.76×10^-9^) and x is the activity of a gene. Dots are colored according to viral lifestyle putative lytic (black) and temperate (gray). Background colors indicate the KEGG metabolisms assigned to the functions. A list of specific genes associated with each pathway or reaction can be found in Supplementary Table S5.

Most pathways investigated were predominantly found on lytic viruses except for other carbon fixation pathways, glycolysis/gluconeogenesis, and the pentose phosphate pathway, which were more often found on temperate viruses (Figure 4B). Only genes related to sulfur metabolisms showed a near 50% split between viral lifestyles (Figure 4B). Lytic viruses carrying metabolic genes related to thiosulfate oxidation and molybdenum cofactor synthesis were more active in Lime Blue, whereas in Poison Lake, lytic viruses carried Vitamin B12 synthesis genes (Tukey’s Test, α < 0.05, Figure 4C). Lime Blue temperate viruses carrying genes related to the pentose phosphate pathway, incorporation of sulfide into cysteine, iron storage, and iron cluster assembly were more active than lytic viruses (Tukey’s Test, α < 0.05, Figure 4C). In Poison Lake, temperate viruses with genes related to the conversion of glycine to heme were more active than lytic viruses, while in Mahoney Lake, temperate viruses with genes involved in the oxidation of sulfate to sulfite were more active than their lytic counterparts (Tukey’s Test, α < 0.05, Figure 4C).

## Discussion

### Decoupling between abundance and activity in phototrophic sulfur bacteria

Understanding the biotic and abiotic factors that control bacterial metabolism and, therefore, the production and expression of biosignatures remains a major challenge with implications for geobiology, global biogeochemistry, and astrobiology. Purple and green sulfur bacteria produce diagnostic pigments and isotopic fractionations that are widely used to reconstruct anoxic photic zones through Earth history, yet the controls on these signatures remain poorly constrained in modern analogs of ancient oceans [5,28,29]. Here, we provide evidence that variability in biosignature expression can arise from a decoupling between bacterial abundance and activity. Across microbial plates in all study lakes, PSB and GSB exhibited contrasting relationships between their relative abundance in the microbial community and their activity (Figure 1B). PSBs consistently showed activity exceeding expectations based on their abundance, whereas GSBs were comparatively less active. Importantly, these trends were consistent across both low- and high-sulfide chemoclines. Together, these observations indicate that sulfide availability alone does not explain the observed decoupling between abundance and activity in PSB and GSB on microbial plates.

### Contrasting viral interactions in PSB and GSB

The elevated activity of PSBs relative to their abundance occurred concurrently with several indicators of lysogenic phage infections, suggesting that lysogeny contributes to sustaining high PSB activity. First, temperate viruses were more abundant and active in the microbial plates of lakes with higher sulfide concentrations when compared to Lime Blue Lake, where sulfide concentration is orders of magnitude lower (Figure 2). Second, all virus-host associations identified for PSBs in the dataset are exclusively with temperate viruses (Figure 3), and the VHRs of abundance and activity of these virus-host pairs were low, indicating limited activity and replication consistent with a temperate lifestyle. Although the high-throughput viral identification methods applied here did not identify a medium or high-quality prophage integrated into the genomes of the dominant PSB genus *Thiohalocapsa*, manual inspection of this genome using VirSorter2 identified two putative prophage regions, each containing at least two viral genes (Supplementary Table S6). Discrepancies between viral prediction tools often arise from differences in databases and models, and the fragmented nature of MAGs can make identifying prophages difficult [75]. Taken together, these observations indicate that PSBs within the microbial plate experience weak top-down viral control and that lysogeny is the predominant viral strategy associated with these populations. Lysogeny could promote higher host activity through a Make-the-Winner regime in which superinfection exclusion increases host fitness [76,77]. Despite the numerical dominance of *Thiohalocapsa* within PSB populations, relatively few viruses were directly linked to this genus, while viral associations were disproportionately observed among less abundant community members (*Bacteroidales*, n=141, 4-14% abundance, 3-5% activity). Similar patterns have been reported in other microbial ecosystems, where viruses have been implicated in controlling populations and promoting diversity [78,79].

In contrast to PSBs, the comparatively low activity of GSB populations coincided with indicators of stronger lytic top-down viral control. The first line of evidence supporting this conclusion is the lower temperature viral abundances in the microbial plate of Lime Blue (dominated by GSB and with trace sulfide concentrations) than in systems with higher sulfide pools (Figure 2). Bacteria containing CRISPR-Cas defense systems were more abundant and active in Lime Blue Lake, including the most dominant GSB, whereas the dominant PSB in lakes with a large sulfide pool only carried RM systems (Figure 2). These results align with the idea that hosts with CRISPR-Cas systems frequently infected with temperate viruses may be lose these systems due to the potential for autoimmunity [80]. Consistent with this pattern, dominant GSB taxa, such as *Chlorobium*, were only associated with lytic viruses (Figure 3), indicating that lytic infection represents the prevailing viral strategy in this system. This interpretation is supported by previous observations showing that total viral abundances and VHRs in the Lime Blue microbial plate, quantified via epifluorescence microscopy, were significantly higher than those in surface waters and in microbial plates from Mahoney and Poison lakes [19]. We speculate that the reduced activity of GSB relative to their abundance could therefore result from lytic infections redirecting host resources and transcription toward virion production and repeatedly removing active cells from the population [42,81–83]. These dynamics contrast with the lysogeny-dominated viral strategies observed in PSB populations, even when both guilds co-occur within the same chemocline. One caveat to the conclusion that GSBs are under lytic top-down control is the relatively low VHRs calculated from abundance and activity of the dominant *Chlorobium* virus-host pair (Figure 3). However, this observation may also be explained by Kill-the-Winner cycles [84], and additional work on this specific pair is necessary to address this open question.

### The infection strategy is host-linked

The consistently high activity relative to abundance of PSBs and their association with temperate viral infections across all three microbial plates with wide ranges of sulfide availability (0.006 – 25.389 mM) suggest that lysogeny is favored by intrinsic host traits rather than by external geochemical conditions. One such trait may be the host’s genome architecture. Lysogeny has been shown to be more prevalent in hosts with larger genomes, which provide increased availability of neutral integration sites, greater regulatory complexity, and enhanced buffering against the metabolic costs of prophage maintenance [85–87]. Consistent with this framework, the average genome size of PSBs within the *Chromatiaceae* (∼4.31 Mb) is approximately 1.75 times larger than that of GSBs within the *Chlorobi* (∼2.47 Mb), potentially increasing PSB susceptibility to stable prophage integration. Differences in antiviral defense strategies further support this interpretation. PSB-associated MAGs with RM systems showed a higher prevalence and activity, whereas MAGs with CRISPR-Cas systems, which typically protect from lytic viruses, were comparatively less abundant and active (Figure 2C–D). The most abundant and active GSB MAGs carried CRISPR-Cas systems, indicating lytic pressure. These contrasting defense architectures could also provide a mechanistic explanation for the observed divergence in viral infection strategies between PSBs and GSBs.

### Sulfate-reducing bacteria and viral regulation of sulfur cycling within microbial plates

Beyond their role in structuring phototrophic populations, viruses appear to modulate sulfur cycling through interactions with SRB [88]. The distribution and activity of viral metabolic genes support a role for viruses in sustaining sulfur cycling within microbial plates, particularly under low sulfide availability. In Lime Blue Lake (0.006 mM sulfide), viruses carrying sulfur-related metabolic genes were disproportionately represented among the most active viruses compared to Mahoney (25.389 mM) and Poison (0.831 mM) Lakes, and associated pathways exhibited higher activity than in the lakes with higher sulfide concentrations (Figure 4). These metabolic genes were primarily linked to sulfate reduction, organic sulfur reduction, thiosulfate oxidation, and sulfur relay systems, indicating a functional emphasis on sulfur regeneration rather than phototrophic metabolism (Figure 4). This pattern contrasts with Mahoney and Poison Lakes, where the aggregated transcriptional activity of viral metabolic genes associated with photosynthesis and glycolysis/gluconeogenesis was significantly higher compared to Lime Blue (Tukey’s Test p < 0.05, Figure 4, Supplementary Table S7). Together, these patterns suggest that in environments where sulfide availability can sustain high rates of anoxygenic photosynthesis, viruses may promote host activity through lysogeny, super-infection exclusion, and phototrophic activity. In contrast, in systems characterized by lower sulfide availability, lytic top-down control may limit GSB activity and favor viral metabolic genes that promote processes that regenerate reduced sulfur (Figure 5).

**Figure 5.**
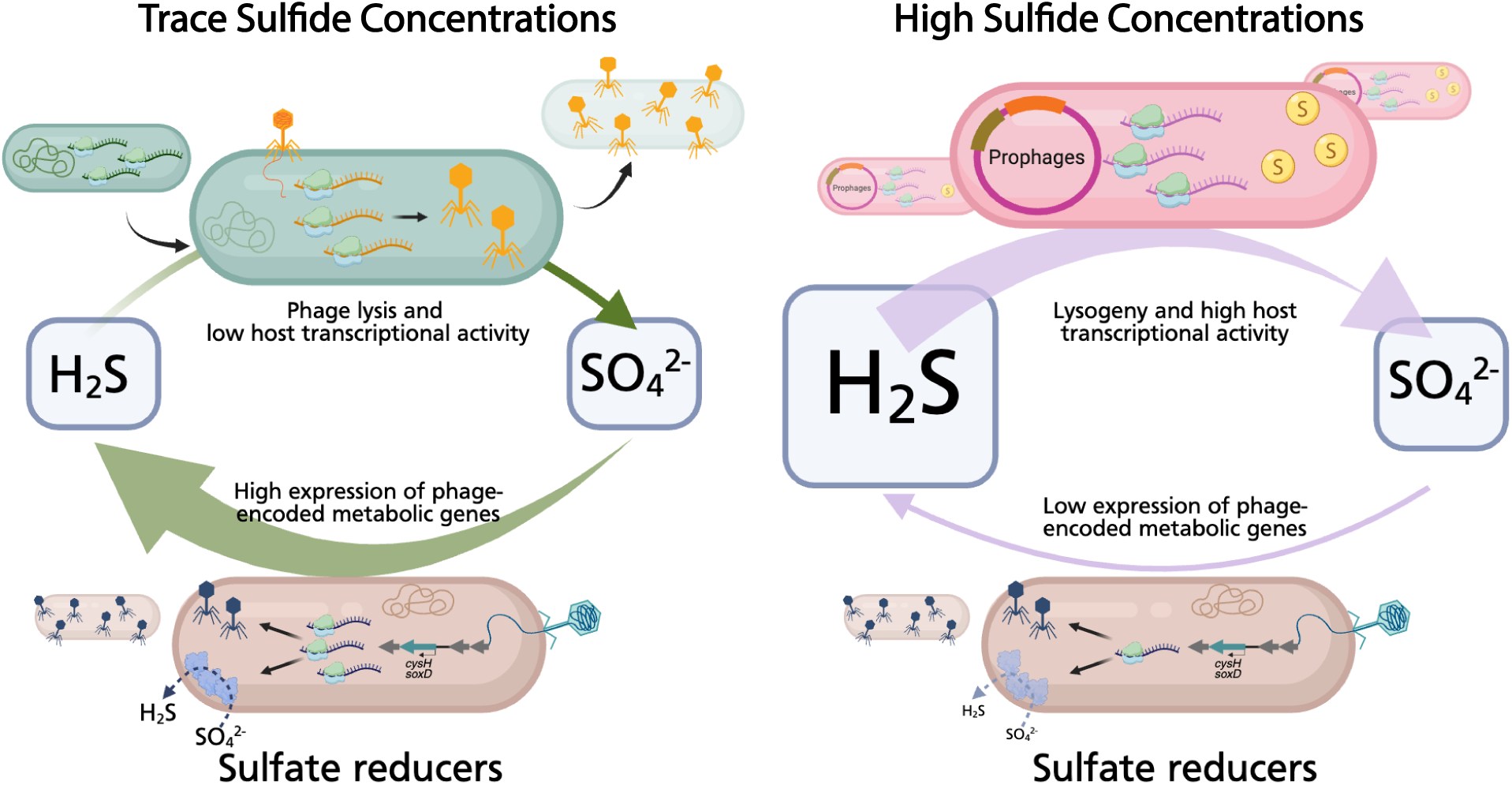
Conceptual framework of contrasting viral controls of phototrophs and sulfur regeneration in meromictic lakes. In systems with trace sulfide-concentrations (left), lytic top-down controls of GSBs reduce host activity (blue mRNA molecules) and redirect transcription to viral propagation (yellow mRNA/viruses), limiting the amount of sulfide oxidized. Viruses carry and transcribe more genes related to sulfide regeneration, helping to sustain the limited pool of sulfide available (bottom, left). In systems with higher sulfide availability (right), integrated viruses (prophages, brown and orange DNA segments) protect the host from lytic infections and promote photosynthetic activity and sulfide oxidation. Activity of SRB (brown cells) and viruses carrying genes related to sulfur regeneration are limited and contributes little to the sulfide pool (bottom, left).

In meromictic lakes, anoxygenic phototrophs are sustained by sulfide supplied both through upward diffusion from the sulfidic monimolimnion and through *in situ* sulfate reduction by SRB occurring within or immediately below the microbial plate [23,30,89]. While sulfate reduction contributes to local sulfide production, the observation that SRB activity is, on average, approximately one order of magnitude lower than that of PSBs and GSBs (Figure 1) suggests that sulfate reduction alone is insufficient to meet phototrophic sulfide demand in such systems.

Within this flux-limited framework, the elevated activity of sulfide-regenerating viral metabolic genes (such as sulfate, thiosulfate, and organic sulfur reduction) in Lime Blue Lake suggests that viral infections may expand the number of pathways that reduce sulfur without substantially increasing overall SRB activity or biomass (Figure 4C). Such viral acceleration of sulfur metabolism could promote rapid sulfur turnover while preventing sulfide accumulation, particularly in systems where sulfide is immediately consumed by highly active phototrophic sulfur bacteria (Figure 5). This tight coupling between sulfate reduction and sulfide oxidation is consistent with models of cryptic sulfur cycling, in which reduced sulfur is continuously regenerated and consumed on short timescales, masking sulfate reduction at the level of bulk geochemistry despite high metabolic activity [23,30,90].

Taken together, these observations indicate that viruses may play a previously underappreciated role in regulating sulfur fluxes within microbial plates by modulating the efficiency and turnover of sulfur cycling pathways. In sulfide-limited systems, such as at the microbial plate in Lime Blue Lake, viral regulation of SRB may therefore contribute to maintaining anoxygenic primary production by buffering sulfur cycling against geochemical limitation.

### Prediction for the geological record

In PSBs, transcriptional activity exceeds abundance, indicating elevated metabolic rates relative to population size. Because PSBs are anoxygenic phototrophs, increased metabolic activity is expected to translate into enhanced photosynthetic rates, which are tightly coupled to light availability [11]. Elevated photosynthesis would promote the synthesis of light-harvesting and storage compounds, including lipid pigments [91–93]. Some of these compounds are recalcitrant, notably the lipid biomarker okenone, which is widely used as a diagnostic indicator of PSB and has been detected in Proterozoic (1.64 billion years) sedimentary rocks [94]. Viral modulation of PSB metabolism could therefore increase the production of PSB-derived biomarkers, increasing the size of the biomarker pool and enhancing the probability that these compounds are preserved in the geological record.

Enhanced metabolic activity may also influence sulfur isotope systematics associated with PSB-dominated sulfur cycling. Higher metabolic rates are generally associated with reduced net isotopic fractionation between substrates and products, as rapid intracellular processing limits isotopic equilibration and isotopic “sorting” between light (^32^S) and heavy (^34^S) isotopes. Microbial sulfide oxidation is generally associated with small sulfur isotopic fractionations and thus often exerts only a limited isotopic overprint on bulk sulfate-sulfide systematics [95]. Notable exceptions exist, for example, sulfide oxidation can in some cases generate substantial ^34^S enrichment in the produced sulfate (+12.5‰) [96]. Under conditions of elevated PSB activity, we therefore predict lower apparent sulfur isotopic fractionation between sulfate and sulfide (Δ^34^S = δ^34^S_sulfate_ − δ^34^S_sulfide_). In this framework, limited top-down control via lytic viral infection could simultaneously enhance the accumulation of PSB-derived biomarkers while dampening the expression of characteristic bulk sulfur isotope fractionation. In contrast, strong lytic viral control of GSBs is expected to reduce overall activity, diminishing pigment production, and weakening the contribution of GSB-derived biomarkers to the sedimentary record.

Beyond phototrophic sulfur bacteria, our observations of elevated activity of sulfate reduction genes in both low- and high-sulfide environments, together with a large number of virus–host associations detected (n=84), suggest that viral infection may also modulate SRB. SRB preferentially reduce ^32^S_sulfate_, and the magnitude of sulfur isotope fractionation expressed during sulfate reduction depends on intracellular sulfate reduction rates as well as extracellular sulfate concentrations^91,92^. In general, sulfur isotope fractionation has a negative relationship with sulfate reduction rates, whereas low intracellular rates can approach near-thermodynamic fractionation even at micromolar sulfate concentrations^93^. The presence of viral metabolic genes linked to sulfate reduction in our dataset raises the possibility that viral infection increases sulfate reduction rates, thereby reducing the net sulfur isotope fractionation between sulfate and sulfide at the bulk scale. Rather than increasing sulfide accumulation, such viral acceleration of sulfate reduction would be expected to increase the sulfide flux through the system, thereby reinforcing rapid sulfur turnover. This enhanced regeneration of reduced sulfur would increase substrate availability for sulfide-oxidizing phototrophs, further coupling reductive and oxidative sulfur cycling within microbial plates.

On early Earth, viral stimulation of sulfate reduction could have contributed to locally or episodically elevated sulfide fluxes in photic zones. Enhanced sulfide availability would have acted as a sink for molecular oxygen and intensified competition for light and nutrients between anoxygenic phototrophs and oxygenic cyanobacteria [1,97]. While viral regulation alone is unlikely to account for global redox states, such biologically mediated feedback could have contributed to maintaining heterogeneity and suppressing net oxygen accumulation at regional scales. Over geological timescales, the cumulative effects of these processes may have contributed to the prolonged delay between the emergence of oxygenic photosynthesis as early as the Eoarchean (∼3.5 Ga) [98,99] and the oxygenation of Earth’s atmosphere (>2.5 Ga) [100], as well as the much later widespread oxygenation of the ocean water column (> 635 Ma) [101,102].

## Conclusion

Within meromictic lakes, microbial plates formed by anoxygenic phototrophs represent hotspots of intense microbial activity and tight biogeochemical coupling. By integrating metagenomics, metatranscriptomics, and virus–host linkage approaches, we show that viral infection strategies in these microbial plates are structured primarily by functional guilds (PSB, GSB, and SRB) rather than by lake-scale geochemical differences. A central outcome of this work is that microbial abundance is not a direct proxy for metabolic activity within microbial plates of anoxic and sulfidic environments. PSBs consistently exhibit activity that exceeds their abundance and are linked exclusively to temperate viruses. In contrast, GSBs display comparatively high abundance but reduced activity and are linked predominantly to lytic viruses. Beyond phototrophs, SRB are associated with both lytic and temperate viruses encoding genes implicated in sulfur transformations, with sulfur-regenerating functions particularly enriched among the active viruses in the sulfide-limited Lime Blue microbial plate.

Our observations indicate that viruses not only act as regulators of microbial population dynamics but mechanistically contribute to how activity is distributed across microbial guilds and how sulfur cycling is sustained within anoxic photic zones. Viral-mediated decoupling of abundance from activity in sulfur cycling bacteria can represent an underappreciated control on the production and the potential for geological expression of sulfur isotope fractionation patterns and lipid biomarker inventories in stratified aquatic environments.

## Acknowledgements

We thank the Frost Institute for Data Science and Computing (IDSC) for providing access to the University of Miami’s high-performance computing system. We thank Alexandra K. Stiffler for laboratory support. We thank Liar’s Cove camping resort for offering power supply and sample refrigeration. We thank Trinity Hamilton for loaning her water-quality sonde and PAR meter. We thank the residents of the Alkali Lake community for providing us with road access to Poison Lake. Mahoney Lake sampling was approved by British Columbia Parks (Permit No. 111692) and facilitated by Wendy Pope, area supervisor of South Okanagan. We thank the Indigenous Peoples of Okanagan for permitting sampling, specifically the SnPink’tn, Osoyoos, and Lower Similkameen Indian Band.

## Data & Code Availability

Transcriptomic and HiC sequence data have been deposited to the National Center for Biotechnology Sequence Read Archive (NCBI SRA) under accession code PRJNA1161448. Metagenomes and MAGs have been published by the Joint Genome Institute and can be accessed using the DOI 10.46936/10.25585/60008509. Detailed description of the bioinformatics pipelines, data tables, and R code for data wrangling and visualization can be found at https://github.com/Silveira-Lab/SPACE2023_lifestyles.

