## Supplementary Figures for "Abundance-activity decoupling in sulfur-cycling bacteria reflects viral infection types in meromictic lakes"

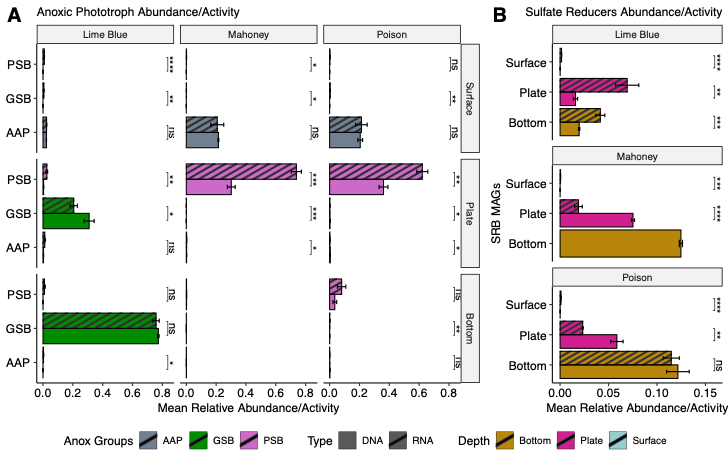

Supplementary Figure 1. Distribution of anoxic phototrophs and sulfate reducers across all samples. A) Relative abundance (solid) and activity (striped) of anoxic phototrophs (A) and sulfate reducers (B). Anoxic phototrophs are grouped as either PSBs (purple), GSBs (green), or aerobic anoxic phototrophs (gray). Sulfate reducers are colored according to the depth of the sample taken surface (blue), plate (pink), and bottom (golden). Results of Tukey’s tests comparing abundance to transcription are denoted by ns = not significant, * = 0.05, ** = 0.01, *** = 0.001, **** = 0.0001.

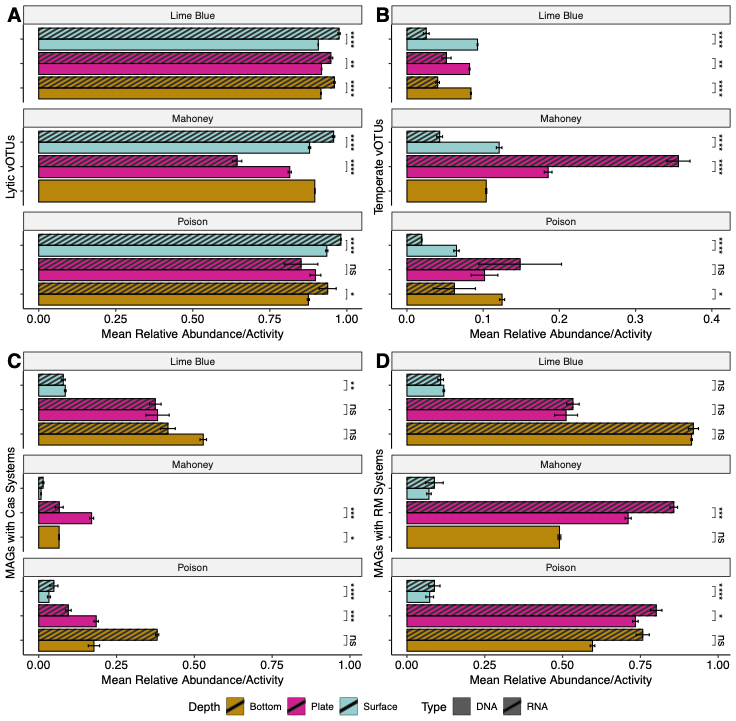

Supplementary Figure 2. Distribution of different viral lifestyles and MAGs containing phage defense system across all samples. Abundance is dented by solid bars and activity striped bar.. Colors of bars represent the layer sampled surface (blue), plate (pink), and bottom (gold). (A) Putatively lytic vOTUs, (B). temperate vOTUs including prophages, (C) MAGs with genes annotated as Cas systems, and (D) MAGs with genes annotated as restriction modification systems. Results of Tukey’s tests comparing abundance to transcription are denoted by ns = not significant, * = 0.05, ** = 0.01, *** = 0.001, **** = 0.0001.

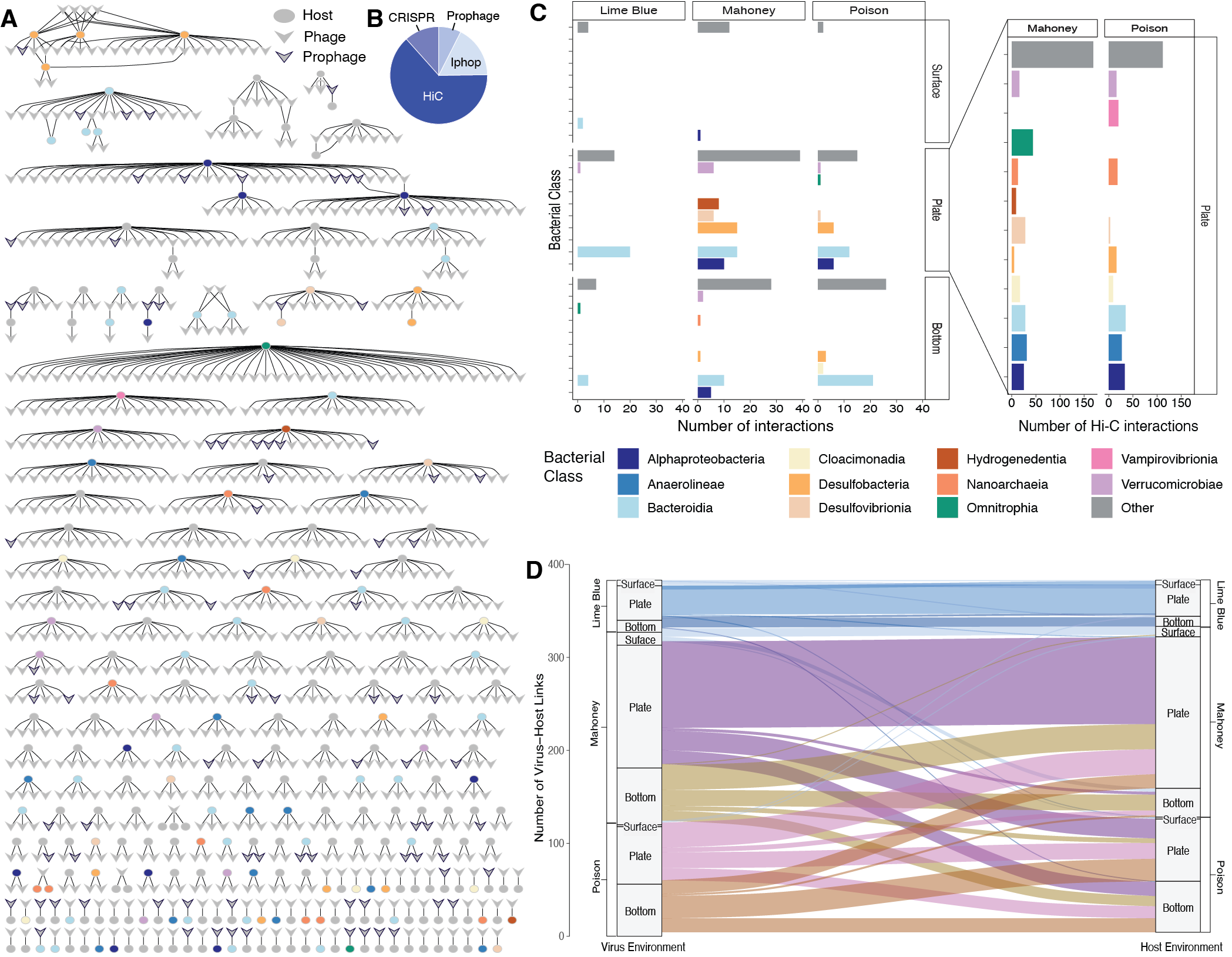

Supplementary Figure 3. Virus-host interaction network. (A) Visualization of interaction between viruses and hosts. Host (circles) are colored according to their class. Phages are indicated by arrows and prophage have a black boarder. (B) Percentage of viruses identified using the four different methods including Hi-C and *in silico* approaches. (C) Distribution of the numbers of connections made by lake and sample depth. Bars are separated and colored according to the host class. The three columns in on the left reflect only *in silico* connections while the 2 columns on the right represent only Hi-C connections. (D) Connections made between original viral contigs (left) assembled from a specific sample type and the sample type that the original MAG (right) was assembled and binned from. Colors indicate the source sample type of the original viral contig.
